# Single-Cell Atlas Uncovers Layer 4 Heterogeneity and Functional Gradients in Rhesus Macaque Visual Cortex

**DOI:** 10.1101/2024.03.11.584345

**Authors:** Dorothee M. Günther, Mykhailo Y. Batiuk, Romain De Oliveira, Viktor Petukhov, Thomas Wunderle, Christian J. Buchholz, Pascal Fries, Konstantin Khodosevich

## Abstract

Non-human primates like rhesus macaques are pivotal models for decoding human visual cortex physiology and disorders. We introduce BrainSPACE, an innovative pipeline for rapid and precise brain tissue banking, to sample visual cortex areas: V1, V2, V4, MT, and TEO. Applying snRNA-seq to V1 and V4 (95,071 nuclei), we uncovered conserved GABAergic neuron profiles but stark area-specific diversity in principal neurons, featuring seven unique layer 4 subtypes in V1 and one in V4. Complementary smFISH validated transcriptional gradients across these areas, aligning with ventral and dorsal stream hierarchies. Gene ontology analyses highlighted plasticity-related pathways in unique layer 4 subtypes, with genes like *NTNG1* and *NLGN1* linked to neurodevelopmental disorders such as autism and schizophrenia. Our insights bridge molecular architecture to visual processing, offering an interactive atlas for community use. By revealing how layer 4 heterogeneity drives hierarchical specialization, our work advances primate brain mapping and informs therapeutic strategies for vision-related pathologies.

## Introduction

Non-human primates (NHPs) such as rhesus macaques are essential for understanding human brain physiology and disorders, particularly visual information processing. Studies in macaques have delineated the visual cortex into specialized areas, including primary visual cortex (V1) and V4, a core model for higher visual processing. V1 sits at the bottom of the visual hierarchy, receiving thalamic input for initial processing^1–3^. From there, information flows to higher areas via two streams: the ventral stream (V1 → V2 → V4 → TEO) performing object recognition^4^, and the dorsal stream (including MT) performing motion and spatial representation^5^.

Thalamic input enters V1 through layer 4, with feedforward projections from V1’s supragranular layers (2/3) targeting V4’s layer 4^3,6^. Feedback from V4 to V1 originates primarily in infragranular layers (5/6)^6^ and avoids V1’s layer 4^3^. Thus, V1 and V4 differ in signal reception, processing, and transmission layers. Yet, how these functional differences manifest in molecular and structural correlates remains unclear. Unlike conserved patterns in mouse V1-V4^7^, macaque exhibits primate-specific divergence, as evidenced in recent studies^8–11^.

Classically, cortical areas are defined by anatomy, layering, neuronal morphology, and connectivity. Recent high-resolution single-cell and spatial transcriptomics studies have revolutionized neuronal diversity mapping, linking it to cortical functions^12–17^. These approaches have generated atlases revealing cytoarchitecture across species^18^, including the rhesus macaque. However, studies comparing neighboring specialized visual areas are scarce, hindering insights into structure-function relationships in visual processing.

Here, we provide a single-cell transcriptomics atlas for early and mid-level visual cortex in the rhesus macaque. We first devised a novel brain tissue banking pipeline for fast and precise sampling, storage and retrieval of rhesus-macaque brain tissue. We implemented the pipeline to identify five areas involved in visual processing, specifically areas V1, V2, V4, MT, and TEO. V1 and V4 were analyzed with snRNA-seq, and all five areas were further analyzed by snRNA-seq-informed multiplexed single-molecule fluorescent in situ hybridization (smFISH). Our snRNA-seq analysis of V1 and V4 revealed high conservation within GABAergic neurons, whereas there was a strong area-specific signature for all principal neuron subtypes. Moreover, seven distinct transcriptomic subtypes were detected specifically among principal neurons of V1, all exhibiting signatures of layer 4, and one subtype specifically among principal neurons of V4 area. Follow-up smFISH analysis for several of these specific subtypes demonstrated transcriptional gradients across V1, V2, V4, MT, and TEO, which appear to largely reflect the functional hierarchy of these areas. Overall, our data show an association between the hierarchical levels and corresponding functions of the investigated areas to their unique organization.

Here, we hypothesize that layer 4 molecular diversity underlies areal specialization. We introduce BrainSPACE for fast and precise tissue sampling and storage, enabling snRNA-seq on V1 and V4, plus snRNA-seq-informed smFISH across V1, V2, V4, MT, and TEO. Our analyses reveal conserved GABAergic neurons but strong area-specific signatures in principal neurons, with unique layer 4 subtypes and transcriptional gradients mirroring functional hierarchies. Community access is provided via interactive platforms.

V1 dataset: (http://kkh.bric.ku.dk/p2auto/index.html?fileURL=http:/rdo/project_1/visualisation_2.bin)

V4 dataset: (http://kkh.bric.ku.dk/p2auto/index.html?fileURL=http:/rdo/project_1/visualisation_3.bin)

Integrated dataset: (http://kkh.bric.ku.dk/p2auto/index.html?fileURL=http:/rdo/project_1/visualisation_1.bin).

## Results

### BrainSPACE - novel pipeline for mammalian brain-tissue banking and storage for multi-purpose use

The rhesus macaque cortex is comprised of roughly 140 anatomically defined areas^19^, and within the visual cortex, 29 areas are distinguished, among them V1, V2, V4, MT, and TEO (Fig. 1A). To enable precise and systematic sampling from banked brain material for follow-up analyses, such as snRNA-seq, we designed a box for dissection and storage, the BrainSPACE (Brain SPatial ArChivE). Within 30 minutes of the onset of cardiac arrest (see Methods for details), the skull was opened (Fig. 1B), and the whole brain was extracted and transferred to ice-cold buffer to reduce the degradation of RNA and proteins in the freshly dissected tissue. We removed the cerebellum and separated the hemispheres along the midline, to be able to position each hemisphere with its relatively flat medial surface on the precooled spacer plate (Fig. 1C). The hemisphere froze onto the spacer plate, attaching the tissue in place and allowing a cut along the spacers into a 2 mm thick sagittal section. Then the rest of the hemisphere was placed, with the freshly cut surface down, onto the next spacer plate and the process was repeated until the whole hemisphere was cut into eight to nine 2 mm thick sagittal sections. All sections were imaged on the spacer plates before storing the spacer plates in the BrainSPACE box at −80°C (Fig. 1B) for further use. Using the images for manual alignment to an atlas^20^, areas were identified and sampled from the respective slices. From the frozen slices, 5-10×10 mm blocks from the respective areas spanning all cortical layers and including white matter were extracted from V1, V2, V4, MT and TEO (Fig. 1 C,D).

**Figure 1:**
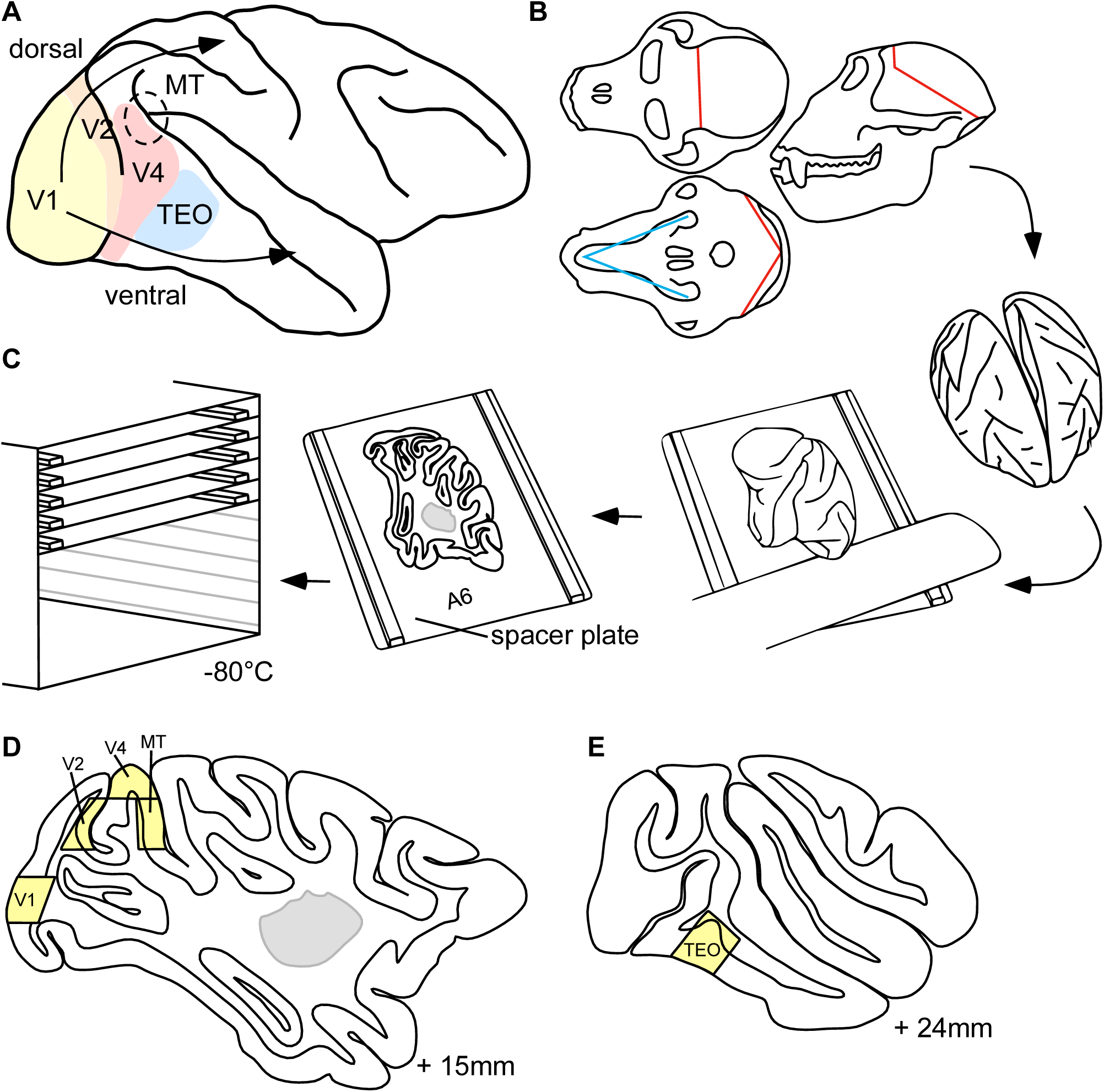
Brain preparation and storage. **(A)** The rhesus macaque brain. Depicted are the primary visual area (V1), visual area 2 (V2), visual area 4 (V4), middle temporal area (MT) and area TEO including their location within the ventral and dorsal stream. **(B)** Drawings of the *Macaca mulatta* skull from top (top left), bottom (bottom left) and side (right). Tissue cuts are marked in blue and bone saw cuts are marked in red. **(C)** Experimental design of the tissue sectioning. Hemispheres were separated, the cerebellum removed, and the hemisphere placed with its medial side onto a cooled plate. The knife was moved along the bars (2 mm height), the rest of the hemisphere removed and placed onto the next plate. The plate was placed on dry ice until frozen and stored in a box at −80°C. **(D)** 15 mm lateral section depicting the extracted tissue blocks for single-cell sequencing and further treatment for smFISH. **(E)** 24 mm lateral section depicting the extracted tissue blocks for single cell sequencing and further treatment for smFISH.

### Taxonomy of neuronal types across visual areas

Tissue samples from areas V1 and V4 from rhesus macaques F354 and M223 (see Methods) were collected and analyzed by single nucleus RNA-sequencing (snRNA-seq) in this study. To this end, 2×5×10 mm tissue blocks were homogenized, and neuronal nuclei were sorted by fluorescent-activated nuclear sorting (FANS) using an antibody to a general neuronal marker NeuN (Fig. 2 A, Supplementary Fig. 1A) as described before^21^. SnRNA-seq was performed using the 10X Genomics 3’ RNA assay. In total, we sequenced 95,071 neuronal nuclei from 2 rhesus macaques (1 male, 1 female), 53,144 from V1 and 41,927 from V4 (4,182 median Unique Molecular Identifiers (UMIs) and 2,164 median genes per nucleus) (Supplementary Fig. 1B-D).

**Figure 2:**
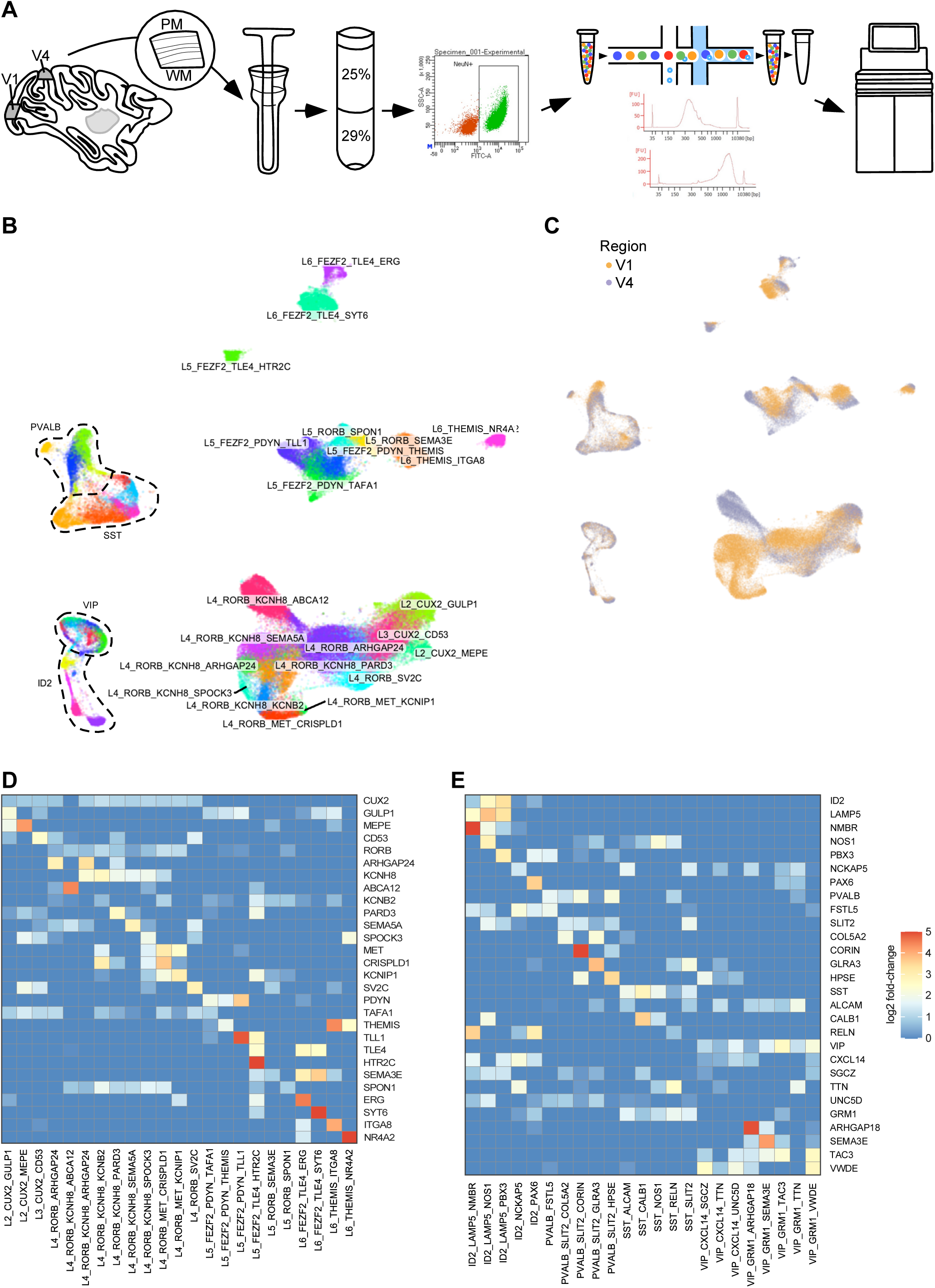
Experimental procedure and snRNA-seq dataset. **(A)** Experimental design of the snRNA-seq measurement: tissue blocks from cortical areas V1 and V4 were dissected from sagittal slices of rhesus macaque brain, tissue was homogenized, nuclei purified using gradient centrifugation and neuronal nuclei were stained using NeuN with following FANS. Neuronal nuclei were sequenced using 10x Genomics assays. **(B)** Major neuronal families and their subtypes were detected using single nuclei RNA sequencing. UMAP representation of single nuclei transcriptomes is shown; nuclei are colored by their subtype. **(C)** UMAP representation of single nuclei, colored by cortical area V1 and V4. **(D)** Heat map depicting expression of markers for principal neuronal subtypes, log2 fold-change is plotted. **(E)** Heat map depicting expression of markers for inhibitory neuronal subtypes, log2 fold-change is plotted.

Neuronal subpopulations were hierarchically annotated across all samples (Fig. 2B). First, to annotate the neuronal populations consistently, inter-individual variations were minimized by aligning the samples from V1 and V4 (Fig. 2C). Further, at the lowest resolution, principal neurons and GABAergic neurons were separated. Followed by the separation of major families of principal and GABAergic neurons using family-specific marker genes for principal neurons (*CUX2*, *RORB*, *FEZF2*, *THEMIS*) and for GABAergic neurons (*ID2*, *PVALB*, *SST*, *VIP*) (Fig. 2 D,E). Next, for each family, we identified fine-resolution transcriptomic subtypes, which in some cases could be grouped in subfamilies based on common markers and similarities present in several subtypes (for instance, subfamilies L5_FEZF2_PDYN and L5/6_FEZF2_TLE4) (Fig. 2D,E). To name subfamily and subtype, we chose among the most specific markers labeling a corresponding cluster, where subtypes with subfamily category had three hierarchical gene markers (family_subfamily_subtype), and those without subfamily category had two hierarchical gene markers (family_subtype). At highest resolution, we could distinguish 46 transcriptomic subtypes, 23 of which belonged to principal neurons and 23 of them to GABAergic neurons (Fig. 2B-E, Supplementary Fig. 2).

To identify potential impact of covariates, such as individual, sex, hemisphere, kit version and sequencing batch, on subtype annotation, we labeled each of these major covariates and showed that they were evenly distributed across UMAPs (Supplementary Fig. 3A-F). We validated clustering robustness by referencing Conos’ built-in batch correction^22^, confirming silhouette-like quality through even covariate distribution (Supplementary Fig. 3A-F).

### Comparative analysis reveals strong areal differences of neuronal complexity in the visual cortex

Integrating V1 and V4 data showed conserved GABAergic expression but pronounced divergence in principal neurons (Fig. 2B and 2C). This contrasts with broader cortical conservation (e.g., mouse/human M1 vs. V1)^7,16^, suggesting unique sub-patterning in macaque visual areas. To cross-validate our findings, we aligned top markers (e.g., *RORB*, *KCNH8*) with two previously published datasets from macaque V1^23,24^ which showed high *RORB* expression in L3-5 (Pearson r inferred ~0.8 for shared markers) and *KCNH8* in L4/5 IT (adjusted P < 1e-300), confirming consistency with our snRNA-seq findings (Supplementary Table 1)^23,24^. Extending to cross-species comparisons, we noted high ortholog conservation (Supplementary Table 1): e.g., *NLGN1* and *SPOCK1* share ~99% identity between human and macaque, with *NTNG1* at 100%, supporting shared synapse formation roles while primate-enriched expression in visual layers diverges^25–28^. Interestingly, most unique L4 subtypes were of the RORB_KCNH8 subfamily (Fig. 3A), which might be a unique subfamily for the visual cortex in rhesus macaque, since strong *KCNH8* gene expression enrichment in the macaque visual vs prefrontal and motor cortices has been reported before^23^.

**Figure 3:**
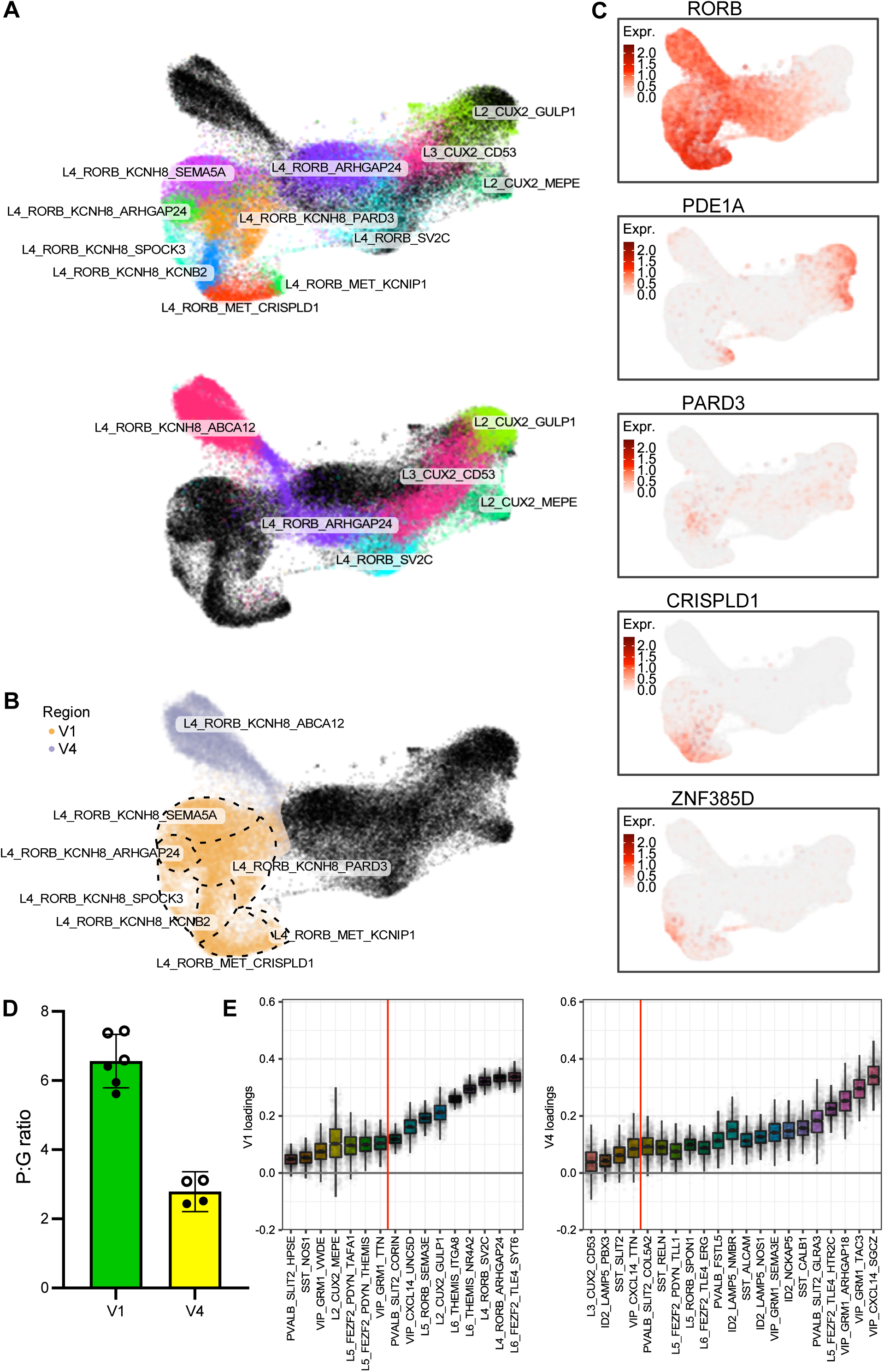
Compositional differences in cell types within visual areas. **(A)** Top plot: V1 layer 4 neuronal subtypes (in color) segregate from V4 subtypes (in grey) mostly by the presence of layer 4 V1-specific neuronal subtypes of L4_RORB family (L4_RORB_KCNH8_SEMA5A, L4_RORB_KCNH8_ARHGAP24, L4_RORB_KCNH8_KCNB2, L4_RORB_KCNH8_PARD3, L4_RORB_KCNH8_SPOCK3, L4_RORB_MET_CRISPLD1, L4_RORB_MET_KCNIP1); Bottom plot: Similarly, V4 layer 4 subtypes (in color) segregate from V1 layer 4 subtypes (in grey) mostly by the presence of V4-specific subtype L4_RORB_KCNH8_ABCA12. UMAP representation of nuclei transcriptomes is shown. **(B)** Expression of marker genes being selected for further smFISH analysis based on their expression in cell clusters uniquely present in V1 layer 4. **(C)** V1 neuronal subtypes (orange) segregate from V4 subtypes (purple) in layer 4. **(D)** Relative number of principal neurons to GABAergic neurons (P:G ratio) in areas V1 and V4. Bar indicated the mean and 95% CI across samples. **(E)** Further change in neuronal composition between V1 and V4 evaluated by compositional data analysis. Cell types distinguishing composition of V1 and V4 samples and present in both regions are shown, region-specific subtypes mentioned above were omitted. The y-axis indicates the separating coefficient for each cell type, with values corresponding to neurons with increased abundance in V1 (top) and V4 (bottom). The boxplots and individual data points show uncertainty based on bootstrap resampling of samples and cells. Red line represents cut-off for significance of adjusted (by the Benjamini-Hochberg method) P values (significant cell types are above the line).

We also noticed a general shift in principal neuron/GABAergic neuron ratio in V1 compared to V4. Thus, V4 had a larger fraction of GABAergic neurons than V1, with the ratio shifted from V1: 6.46 to V4: 2.75 (Fig. 3D; Supplementary Table 1). Compositional analysis revealed V1 enrichment in principal subtypes (mainly L4-6) and V4 in GABAergic subtypes (especially VIP) (Fig. 3E; p<0.05, Benjamini-Hochberg adjusted). One of the most striking changes was in L4_RORB_KCNH8_PARD3 that exhibited large expansion in V1 as 25.3% ± 4.1%, whereas in V4 it accounted only for 8.2% ± 2.3% (p<0.001; Supplementary Table 2). It was opposite for V4, where increases in GABAergic neuron subtypes were the most significant, with VIP neuron subtypes exhibiting the largest enrichment (Fig. 3E). This indicates potential differences in excitation/inhibition regulation in V1 and V4, where V1 has increase in excitatory neurons in middle-deep layers, whereas V4 has increase in VIP neurons that usually disinhibit excitatory neurons by inhibiting PVALB/SST GABAergic neurons.

### Comparative Functional and Molecular Profiling of Unique and Shared Principal Neuron Subtypes in V1 and V4

To explore pathways that might shed light on the function of unique neuronal subtypes in V1 and V4, we extracted the top up- and down-regulated genes for each neuronal subtype and calculated the enriched Gene Ontology (GO) terms. For up-regulated enriched GO terms in neuronal subtypes unique to V1 and V4, we revealed major clustering of GO terms (Fig. 4A). Strikingly, all eight unique principal neuron subtypes in layer 4 of V1 or V4 formed a cluster with enriched GO terms related to cell adhesion and neuron development (red box, Fig. 4A), which might indicate enhanced plasticity of these unique neurons. Analysis of top down-regulated GO terms for unique neuronal subtypes showed subtype-specific enrichment of GO terms, mainly involved in ion transport, cell signaling and synaptic processes (Supplementary Fig. 5A).

**Figure 4:**
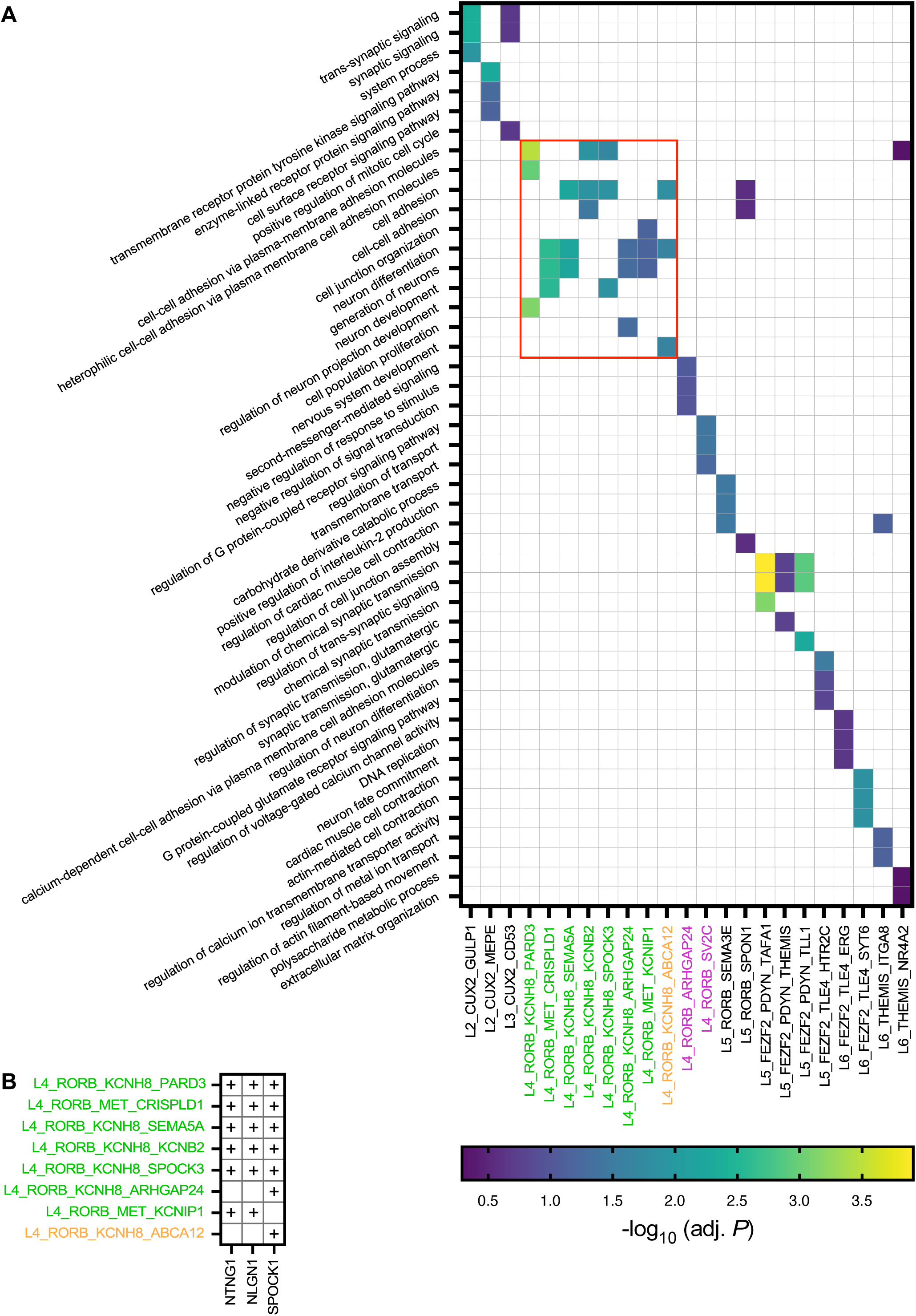
Enriched GO terms show similarities between unique V1 and V4 subtypes. **(A)** Heat map of the top three enriched GO terms for each annotated cell type, colored by – log₁₀(adjusted p-value). GO term enrichment was calculated using a hypergeometric test and adjusted for multiple comparisons by the Benjamini–Hochberg procedure. Cell types unique to V1 are in green, unique to V4 in orange, and layer 4 types shared between V1 and V4 in purple. A red box was manually added to delineate a notable cluster encompassing all eight unique layer 4 neuronal subtypes, highlighting their shared enrichment in GO terms related to cell adhesion and neuron development (see main text for details). **(B)** Dot plot of genes annotated to the top three enriched GO terms for V1-unique (green) and V4-unique (orange) cell types, with dot size proportional to –log₁₀(adjusted p-value) and color indicating average log₂ fold-change.

Interestingly, two common subtypes found in layer 4 of both V1 and V4 (L4_RORB_ARHGAP24, L4_RORB_SV2C), although belonging to the same family of RORB neurons, were not part of the red box cluster and did not show enrichment in cell adhesion and neuron development GO terms (Fig. 4A). Analysis of the genes associated with the enriched GO terms showed frequent occurrence of the genes *NTNG1*, *NLGN1*, and *SPOCK1*. While *NTNG* and *NLGN1* appear to be highly expressed in the same cell types in V1, *SPOCK1* is highly expressed in L4_RORB_KCNH8_ABCA12 (V4-specific) and L4_RORB_KCNH8_ARHGAP24 (Fig. 4B). *NTNG1* is known for its role in neurite outgrowth^29^ and is associated with Rett syndrome^30^ and schizophrenia^31^. *NLGN1* is associated with synapse formation^32^ and autism^33–35^, while *SPOCK1* is associated in neural development^36,37^.

### Gradient principle of L4 neuronal subtypes distribution in the areas of the visual cortex

To reveal general principles of distribution of V1/V4-unique subtypes of L4 principal neurons across visual cortex areas, we set focus on the layer 4 principal neuron subtypes. We selected four V1-unique subtypes for smFISH analyses: L4_RORB_KCNH8_PARD3 (PARD3), L4_RORB_KCNH8_SPOCK3 (ZNF385D), L4_RORB_MET_CRISPLD1 (CRISPLD1), L4_RORB_MET_KCNIP1 (PDE1A) (Fig. 3B) and designed probes targeting *CAMK2A* (principal neuronal marker), *RORB* (neuronal marker, expression in L2/3-5, peak expression in L4^7,38^), and subtype specific markers^39,40^ (Fig. 5A). Upon labeling, layers were delineated based on the density of nuclei y (DAPI), Nissl staining, as well as CAMK2A and RORB expression (Fig. 5B) followed by segmentation and quantification of expression that assigned subtypes via marker co-expression (Fig. 5C).

**Figure 5:**
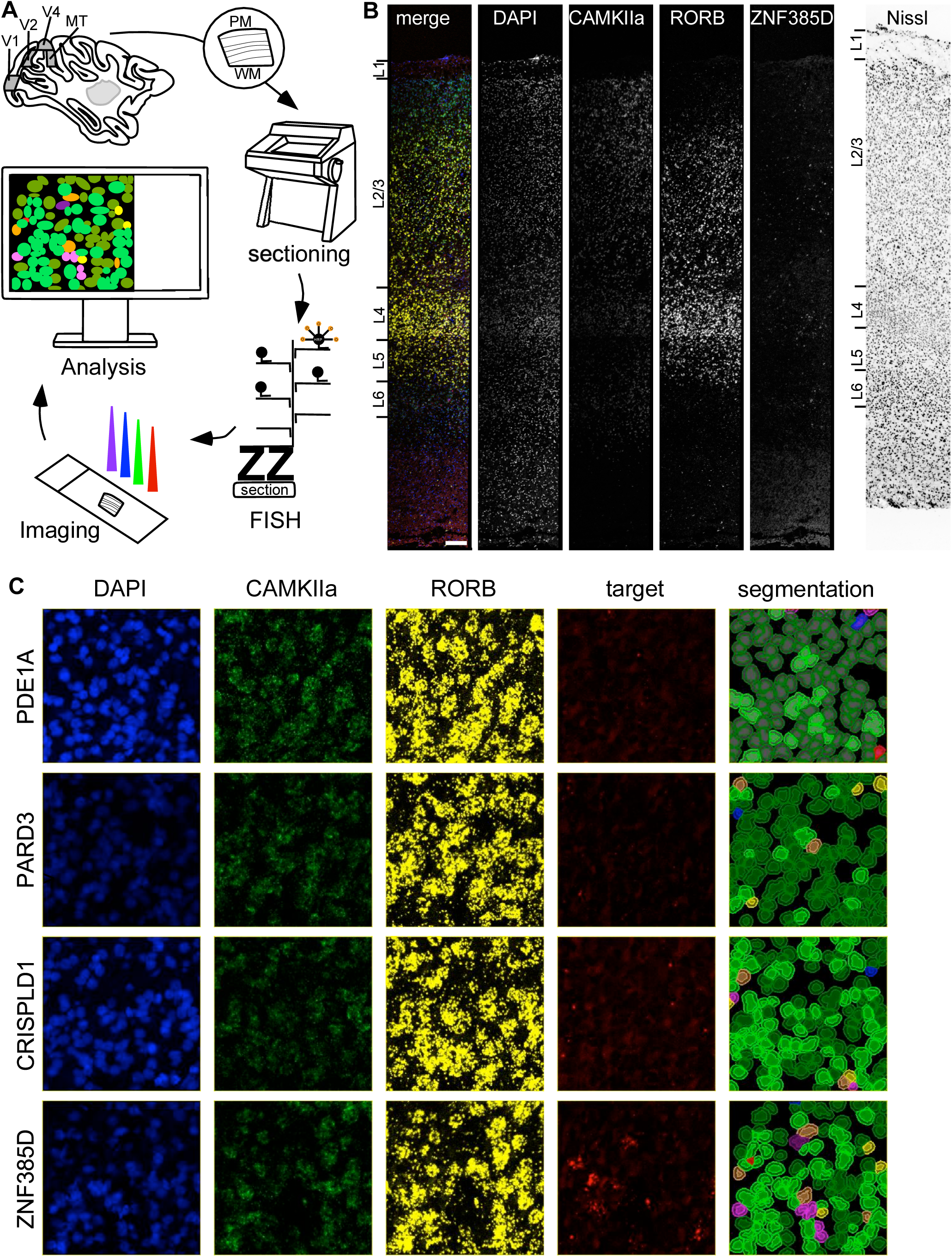
Experimental design for quantitative smFISH. **(A)** Experimental design of the fluorescent in situ hybridization workflow. Tissue blocks were extracted from sagittal brain slices, sectioned into 10 µm sections, conducted to the RNAscope Multiplex Fluorescent V2 Assay, visualized utilizing a Leica SP8 confocal microscope and analyzed using the QuPath software. **(B)** Overview image showing an exemplary staining. Laminar patters for *CAMK2A*, *RORB*, the respective target and Nissl from the pia mater to the white matter. Fluorescent imaged from fourfold stain 10 µm thick section from V1. Scale bar: 200 µm. Brightfield image from 10 µm section from V1. Scale bar: 200 µm. **(C)** Exemplary single-color images from layer 4 in V1. DAPI nucleus staining: blue; *CAMK2A* hybridization: green; *RORB* hybridization: yellow and target hybridization: red. Segmentation: pink: negative; red: target +; blue: *CAMK2A* +; yellow: *RORB* +; dark green: *CAMK2A* + *RORB* +; orange: *RORB* + target+; purple: *CAMK2A* + target+; light green: *CAMK2A* + *RORB* + target+. Scale bar: 45 µm.

Quantification showed inter-individual variability (1.5-40% positive cells; up to 65% in individual sampling windows), reflecting biological heterogeneity (coefficient of variation (CV) ~47.5%; Fig. 6A-D). High positivity for single replicates indicates that localized regions or specific counting windows may show higher enrichment of certain neuronal subtypes, possibly reflecting true biological hotspots (secluded area with cells showing higher gene expression than in surrounding cells) or variation in probe accessibility and signal detection across different tissue sections^41^. Three subtypes - PARD3, ZNF385D, CRISPLD1 were enriched in V1 vs. V4 (p<0.001, chi-square, Bonferroni-corrected) with decreasing abundance along the hierarchy V1 > V2 > V4>MT; V4≤TEO (Fig. 6B-F; Table 2 for layer-wise statistical comparisons). The fourth subtype - PDE1A - showed an opposite gradient (V1 < V2 (peak ~35%)>V4<MT<TEO) (Fig. 6A, E, F).

**Figure 6:**
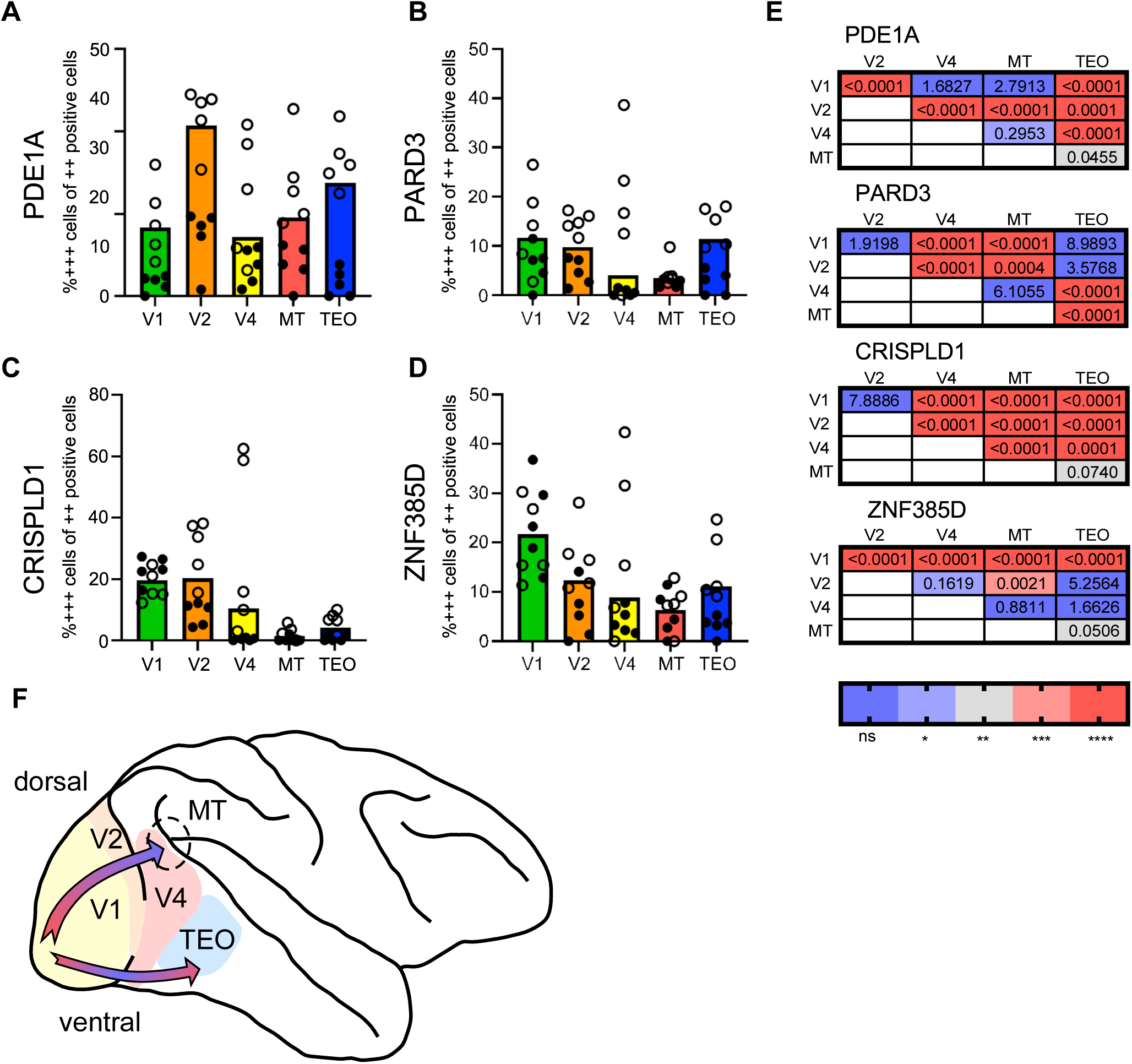
Gradient of unique subtypes of principal neurons along dorsal stream in visual cortex areas. **(A-D)** Percentage of *CAMK2A*+ *ROR*B+ target+ cells of *CAMK2A*+ *RORB*+ cells in layer 4 over V1, V2, V4, MT, and TEO. Symbols represent data from 10 counting windows over n = 2 rhesus macaques, circles represent monkey E and dots represent monkey S, bar represent means, error bars represent 95% CIs. **(E)** Significance (single cells chi-square tested, Bonferroni corrected). **(F)** Model showing the gradient development along the dorsal and ventral stream. Density of cell typed decrease along the dorsal stream while along the ventral stream density of cell types decrease until V4 but increase in TEO again.

Layer-wise, PARD3, ZNF385D, CRISPLD1 subtypes peaked in V1 layer 4 (decreasing in 2/3 and 5), while for V2, V4, MT, and TEO, the expression was generally lowest in layer 4, further confirming that these markers are most strongly associated with the highly specialized layer 4 in V1. For PDE1A+ the expression was lowest in layer 2/3 and peaked in layer 5 (Fig. 7A-E). Overall, these results highlight the unique and shared characteristics of layer 4 in visual areas, reflecting areal specialization for visual information processing.

**Figure 7:**
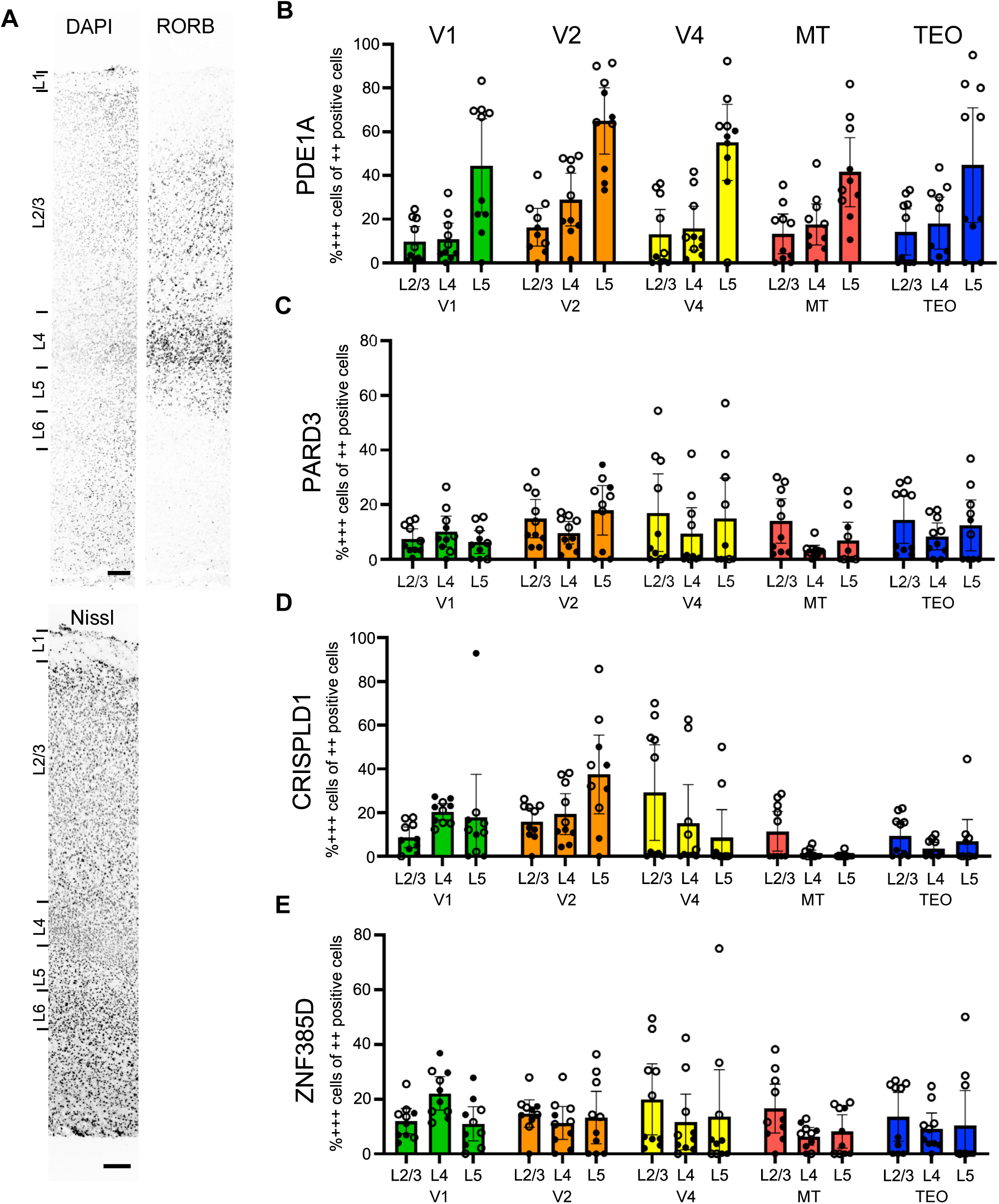
Enrichment of unique subtypes of principal neurons in layer 4. **(A)** Overview image showing an exemplary staining. Laminar patters for DAPI, *RORB* and Nissl from the pia mater to the white matter. Fluorescent images (DAPI and RORB) and brightfield image (Nissl) from V1. Scale bar: 200 µm. **(B-E)** Percentage of *CAMK2A*+ *RORB*+ target+ cells of *CAMK2A*+ *RORB*+ cells across layer 2/3, layer 4 and layer 5 for V1, V2, V4, MT and TEO. Symbols represent data from 10 counting windows over n = 2 rhesus macaques, circles represent monkey E and dots represent monkey S, bar represent means, error bars represent 95% CIs.

## Discussion

The rhesus macaque brain, closely related to the human system, has proven invaluable for elucidating visual processing. Our analysis of areas V1, V2, V4, MT, and TEO reveals substantial cellular complexity that mirrors their hierarchical organization. V1 is the most extensively studied visual area, while V4 serves as a primary model for higher ventral-stream processing. By sampling V1 and V4 separately for detailed transcriptomic comparison, we uncover anticipated differences grounded in anatomy and function. V1, also known as striate cortex, features the prominent stria of Gennari, a band of heavily myelinated axons within layer 4 sublayers, that is visible even to the naked eye in unfixed human brain tissue. Extensive research has delineated distinct sublayers in V1 layer 4 with differential connectivity patterns^39^, including a high proportion of spiny stellate cells^40^. These features likely underpin its role as the main cortical entry point for thalamic (LGN) visual inputs.

To facilitate investigations of banked brain material for follow-up analyses, we developed BrainSPACE, a novel pipeline for precise sampling and storage of unfixed macaque brain tissue, supporting multi-modal analyses. We applied this approach to ~100,000 neurons via snRNA-seq from V1 and V4, identifying 23 GABAergic and 23 principal neuron subtypes and assigning the latter to cortical layers based on marker genes. Our atlas uncovers layer 4 as a hub of areal specialization in primate visual cortex, with gradients mirroring visual information processing hierarchies. Building on recent primate single-cell transcriptomics studies^8–11,13,42–44^, we fill gaps in visual areal organization, linking subtypes to functional streams. While conserved GABAergic neurons align with prior reports^45^, principal neuron divergence, especially layer 4 heterogeneity, highlights visual-specific adaptations, potentially driven by genes like *KCNH8*^46^ and *RORB*^38,47–49^.

GABAergic subtypes show robust conservation between V1 and V4, aligning with patterns across primates^7,13,42–44^. For instance, PVALB+ GABAergic neurons are enriched in the occipital lobe (i.e. visual areas) of marmosets^44,45^, macaques^43^ and humans^16^. Interestingly, few subtypes of GABAergic neurons, including those from SST, PVALB, and VIP families, were reported to be specific to human V1^16^ and enriched in macaque occipital/parietal regions^43^. Our sampling from two occipital areas yielded nearly identical GABAergic profiles, thus demonstrating that such V1-specific subtypes of GABAergic neurons are likely extended to other areas of visual cortex. In contrast, principal neurons in general display marked area-specific signatures – while subtypes are generally shared across cortical regions in varying proportions^16,38,43,44^, pronounced differences emerge even between adjacent areas, as seen in human cortex^16^. Here we also confirm this specialization for macaque visual cortex. However, within visual cortical areas, we identified specific subtypes of principal neurons only in layer 4, indicating high divergence of this layer, likely due to functional specialization of each visual cortex area. We also confirm a higher principal-to-GABAergic ratio in V1 than V4^16,43^, highlighting excitation-inhibition shifts. The shift of principal-to-GABAergic neurons ratio suggests excitation-inhibition balance differences between visual areas. Thus, V1’s enrichment in excitatory neurons may support granular input processing, high-resolution processing of initial thalamic inputs, whereas V4’s increased proportion of GABAergic VIP neurons could enable disinhibition mechanisms, since VIP neurons often inhibit SST and PVALB neurons^50^, thereby releasing principal neurons from suppression and facilitating the contextual integration of complex visual features at higher hierarchical levels.

One of our key findings highlights V1’s distinctiveness: seven “transcriptomically” unique layer 4 principal neuron subtypes in V1 (absent in neighboring V4), which reinforces the specialized function of V1 layer 4^40^. Comparable V1-specific layer 4 subtypes have been reported in marmosets, macaques, and humans^13,43,44^. All seven subtypes express the family marker *RORB*, observed in V1 layer 4 of cynomolgus macaques^43^ and marmosets^44^, as well as in primary areas of other sensory systems (e.g., somatosensory^47^). Five subtypes also express *KCNH8*, a marker enriched in marmoset visual layer 4^44^. GO analysis associates these subtypes with the *NTNG1* gene, consistent with prior reports^43^. Enriched pathways in unique layer 4 subtypes of principal neurons imply their increased plasticity, with disease-linked genes (e.g., *NTNG1*, *NLGN1*) offering translational insights for autism and schizophrenia^25^.

We extended snRNA-seq findings using quantitative smFISH across V1, V2, V4, MT, and TEO, uncovering density gradients along visual streams for four unique subtypes of principal neurons: L4_RORB_MET_CRISPLD1 (marker: CRISPLD1), L4_RORB_KCNH8_SPOCK3 (marker: ZNF385D), L4_RORB_KCNH8_PARD3 (marker: PARD3), and L4_RORB_MET_KCNIP1 (marker – PDE1A). CRISPLD1, ZNF385D, and PARD3 labeling show similar decline for corresponding subtypes in density dorsally from V1 to MT, paralleling rostral-caudal^16,38,48^ and hierarchical^43^ gradients. Their ventral labeling shows a decrease to V4 and stabilizing at TEO. smFISH confirms primarily enrichment of these three subtypes in V1 layer 4, with diminished presence in layers 2/3 and 5. Markers CRISPLD1, ZNF385D, and PARD3 exhibit hierarchical and laminar decreases, whereas PDE1A rises toward higher areas (peaking in V2) and layer 5, concordant with bulk sequencing data^49^. Observed discrepancies, such as sparse detection of V1-specific subtypes in V4 layer 4 via smFISH but their absence in snRNA-seq, primarily arise from methodological differences: smFISH uses only three markers for spatial resolution, while snRNA-seq leverages genome-wide data without spatial context. Additional factors include nuclear-restricted RNA in snRNA-seq and its relatively lower sequencing depth^43^.

Our BrainSPACE pipeline and interactive atlas enhance reproducibility and community exploration, advancing structure-function studies in primate vision and informing therapies for cortical disorders. Although one of our limitations is small sample size (n=4 macaques, Supplementary Table 3), inter-individual CV ~47.5% supports robustness of our data, consistent with macaque variability studies.

Recent primate single-cell atlases demonstrate the power of single-cell and spatial omics technologies for brain complexity^13,42–44^. By resolving visual cortex areal organization, we fill key gaps, aiding insights into visual processing. Our datasets, available through interactive web tools for V1, V4, and integrated datasets, provide a valuable resource for comparative studies and subtype-specific targeting, such as via differentially expressed genes. Since structure-function studies linking neuronal subtypes to their functional roles remain a critical challenge, our atlas will help guide future work to integrate electrophysiology, optogenetics, and connectivity tracing to functional roles for individual neuronal subtypes.

## Materials and Methods

### Rhesus macaque tissue preparation

Rhesus macaque handling and housing conditions were in accordance with the German law for the protection of animals and the “European Union’s Directive 2010/63/EU” and approved by the local government office (Regierungspräsidum Darmstadt).

Rhesus macaques E and S were euthanized due to medial indications at high age. Brains from rhesus macaques 223 and 354 were kindly provided by the veterinary department of the Paul-Ehrlich-Institut, after they had been euthanized for collecting hematopoietic organs for another study (Supplementary Table 3).

Rhesus macaque brain tissue was obtained from 4 *Macaca mulatta* in total, 2 females and 2 males (Supplementary Table 3). As soon as death was confirmed by the veterinarian, the spine was broken and the tongue removed by two incisions (Fig. 1B – blue line). The head was cut off and the skull was exposed by pulling the skin and muscles from posterior to anterior up to the frontal skull. A hand saw was used to carefully open the skull along the red lines indicated in Fig. 1B. The calotte was removed using a spatula, and the brain was pulled out of the skull by gravity. Cranial nerves were cut, and the brain was washed in ice-cold saline. The cerebellum was removed by inserting a spatula from posterior in between the occipital lobe and the cerebellum moving it into anterior direction until the stem brain was separated. The hemispheres were separated along the midline using a Virchow brain knife. The BrainSPACE box, made from stainless steel, was precooled to −80°C over night. Before cutting the brain, the spacer plates with 2 mm space bars on two of their edges were placed onto dry ice. For the process of cutting, a spacer plate was removed from the dry ice and handled at room temperature. The hemisphere was placed onto the pre-frozen spacer plate with the medial side facing the plate. We waited until the tissue was frozen to the spacer plate indicated by tissue color change. Before the frozen tissue part reached the limit of the space bar, the brain knife was used to cut along the spacers, resulting in 2 mm thick tissue sections. Each brain slice was imaged using a high-resolution digital camera. The plate carrying the brain slice was placed on dry ice for complete freezing, and the remaining hemisphere was placed, with the freshly cut surface down, onto the next pre-cooled plate. This sequence was repeated until one entire hemisphere was cut into slices and then repeated for the second hemisphere. Plates were stored in the BrainSPACE box at −80 °C. Each hemisphere resulted in eight to nine sections depending on the size of the animal’s brain.

For *in situ* hybridization, images obtained from the sagittal slices were compared to the rhesus macaque brain atlas^20^, and areas V1, V2, V4, MT and TEO were identified according to the macroscopic anatomic landmarks like sulci and gyri. At the desired locations, the 2 mm thick slices were further cut to obtain 5 × 10 mm tissue cubes, using a hot scalpel as depicted in Fig. 1C. Blocs were embedded in Tissue Freezing Medium (Leica, 14020108926) and stored at −20°C until further processing.

### Nuclei isolation

Nucleus isolation was done as described before with slight modifications^22^. Tissue was removed from the −80°C and transferred into a 1 ml Dounce homogenizer holding 1 ml of prechilled homogenization buffer (250 mM Sucrose, 25 mM KCl, 5 mM MgCl2, 10 mM Tris pH 8.0, 1 mM DTT, 1 x protease inhibitor (Roche, 11873580001), 0.4 U/ul RNAse inhibitor (Takara, 2313B), 0.2 U/ul SUPERase•In (Invitrogen, AM2696), 0.1 % Triton X-100). Tissue was treated with 5 strokes of the loose pestle, followed by 15 strokes of the tight pestle and then filtered through a 40 µm cell strainer. Nuclei were spun down at 1000 × g for 8 min at 4 °C. Supernatant was aspirated, homogenization buffer was added to a total volume of 250 µl and nuclei were resuspended. Nuclei suspension was mixed with 250 µl 50 % Iodixanol (60 % stock from STEMCELL Technologies, 7820), 1 x protease inhibitor (Roche, 11873580001), RNAse inhibitor (0.4 U/ul; Takara, 2313B), SUPERase•In (0.2 U/ul; Invitrogene, AM2696) and 1 mM DTT and layered on top of a 29 % Iadixanol solution (25 mM KCl, 5 mM MgCl2, 10 mM Tris pH 8.0, 29 % Iodixanol, 1 x protease inhibitor, 0.4 U/ul RNAse inhibitor, 0.2 U/ul Superasin, 1 mM DTT) into an ultracentrifugation tube on ice. Gradients were placed into a swing bucket rotor (Beckman Coulter, TLS 55) and centrifuged at ~14.000 × g max (~10900 × g average) for 22 min at 4 °C (Beckman Coulter, MAX-XP). After centrifugation, supernatant was removed, nuclei were resuspendend in BSA blocking buffer (1 x PBS, 0.5 % BSA, 1 mM DTT, 2.54 mM MgCl2, 0.2 U/ul RNAse inhibitor) and incubated for 15 min on ice. Before staining, splits were taken for the controls (isotype control, negative control, 7AAD only control, NeuN only control). To the sample and NeuN only control, the neuronal marker NeuN antibody was added (1 µg/µl, 1:16949, Millipore, MAB3777X). To the isotype control, the control antibody was added (0.2 µg/µl, 1:3390, STEMCELL Technologies, 60070AD). Antibodies were incubated for 10 min at 4 °C in the dark. After the incubation, 1 ml BSA blocking buffer was added and centrifuged at 1000 × g for 10 min at 4 °C in a swing bucket. Pellets were resuspended in 200 µl BSA blocking buffer and filtered through a 35 µm strainer. Samples were filled with BSA blocking buffer to a total volume of 500 µl, and 0.75 µl of 7AAD (1 mg/ml, Sigma) was added. FACS was performed using BD FACSAria III sorter using a 75 µm nozzle and controlled by BD FACSDiva 8.0.1 software. Single color controls were used for compensation. Nuclei were sorted based on the gating strategy depicted in Supp. Fig 2A. Nuclei were collected into 5 µl of BSA blocking buffer at 4 °C. Nuclei were directly processed for 10x library preparation.

### 10x library preparation and sequencing

Chromium Single Cell 3’ Reagent kit v2 (10x Genomics, 120237) and v3 (10x Genomics, 1000075) were utilized for library preparation. The standard protocol was applied as described before^21,22^. In brief, nuclei were counted under a brightfield microscope and mixed with the reverse transcription mix. Either v2 or v3 Gel Beads were added, and the mix was partitioned on Chromium Chips A (v2 kit, 10x Genomics, 120236), or Chips B (10x Genomics, 1000073) into GEMs using the Chromium Controller (10x Genomics, PN-120223), and reverse transcription was performed. Then, samples were frozen at −20 °C until further processing. cDNA was cleaned up, pre-amplified (12 PCR cycles), SPRIselect cleaned up and quantified before being frozen again at −20 °C. If possible, the same amount of cDNA was used for fragmentation, end-repair and A-tailing. The following process was performed: SPRIselect clean up, adapter ligation, SPRIselect clean up, sample index PCR using Chromium i7 sample indices (10-13 PCR cycles, depending on cDNA input quantity, 10x Genomics, PN-120262), SPRIselect clean up and quantification using the Qubit HS dsDNA Assay kit (Thermo Fisher Scientific, Q32854) on the Qubit Fluorometer and the High Sensitivity DNA Kit (Agilent, 5067-4626) on the Bioanalyzer. The number of nuclei was estimated and the libraries pooled accordingly. Pools were again quantified and sequenced on the Illumina NextSeq500 using NextSeq500/550High Output Kit v2.5 (Illumina, 20024907). The sequencing protocol included 26 cycles for read 1, 8 cycles for i7 index and 98 cycles for read 2 for 10x v2 kit or 28 cycles for read 1, 8 cycles for i7 index and 91 cycles for read 2 for 10x v3 kit.

### Single nuclei sequencing data analysis

Primary data analysis was performed using the 10X Genomics CellRanger toolkit (version 3.1.0; https://support.10xgenomics.com/single-cell-gene-expression/software/pipelines/latest/what-is-cell-ranger). The reference genome sequence and annotation used were from Release 98 of the rhesus macaque genome from Ensembl (http://ftp.ensembl.org/pub/release-98/fasta/macaca_mulatta/dna/Macaca_mulatta.Mmul_10.dna.toplevel.fa.gz; http://ftp.ensembl.org/pub/release-98/gtf/macaca_mulatta/Macaca_mulatta.Mmul_10.98.gtf.gz). The annotation file was further filtered to generate a GTF file for creating a pre-mRNA reference, where “transcript” was renamed to “exon” and retained in the final GTF using the following command: “awk ‘BEGIN{FS=“\t”; OFS=“\t”;} $3 == “transcript”{$3=“exon”; print;}’ Macaca_mulatta.Mmul_10.98.gtf > Macaca_mulatta.Mmul_10.98_PRE_mRNA.gtf’. Subsequently, the reference genome was generated for subsequent analyses using “cellranger mkref.”

The Illumina “.bcl” files generated were initially demultiplexed using the “cellranger mkfastq” pipeline. Subsequently, reads were mapped against the reference genome, and UMIs were counted using the “cellranger count” pipeline. Low quality nuclei and empty drops were filtered out using cutoff on feature-count plot.

### Dimensionality Reduction and Data Integration

To visualize the structure and relationships within our integrated dataset, we applied the Uniform Manifold Approximation and Projection (UMAP) algorithm implemented in the conos package^51^ (version 1.5.1). This dimensionality reduction step facilitated the joint embedding of cells from all samples, allowing for the identification of shared and unique cell type clusters across experimental conditions and areas. Subtype markers were then identified using the Conos “getDifferentialGenes” function (version 1.5.1) and further analyzed by manually inspecting marker expression patterns over UMAP maps of subtypes.

### Compositional Data Analysis

We assessed differences in cell-type proportions across samples and conditions using the Cacoa R package^52^ (v0.4.0; https://github.com/kharchenkolab/cacoa) which provides a full R-based re-implementation of the Dirichlet-multinomial compositional testing framework originally described in^22^. This re-implementation preserves the core statistical model and permutation-based significance testing but extends scalability to tens of thousands of cells via sparse-matrix support, incorporates additive log-ratio regression for built-in adjustment of co-variates (e.g. batch, sex), automates Benjamini–Hochberg false-discovery-rate correction across all cell-type contrasts, and integrates streamlined plotting and robustness assessments. Follow-up applications of Cacoa have demonstrated its stability and reproducibility across diverse case–control single-cell RNA-seq studies^53–57^.

We conducted post-hoc power analyses on existing data to confirm statistical robustness. For chi-square comparisons in smFISH (e.g., PARD3+ subtype enrichment between V1 and V4; N=40 windows across areas), we assumed a medium-large effect size (w=0.5) and alpha=0.001, yielding ~45% power. This conservative estimate enables reliable detection of significant differences (p< 0.01), underscoring the dataset’s sensitivity for hypothesis generation while highlighting the need for larger cohorts in follow-up studies^28^. To quantify inter-individual variability (e.g., positivity 1.5– 40%, with hotspots up to 65%), we calculated the coefficient of variation (CV) as ~47.5%, consistent with expected biological heterogeneity in primate samples without undermining core findings. Analyses used the stats model libraries in Python (v3.12.3) for chi-square tests and NumPy^58^ (v2.3.0). All p-values were Benjamini-Hochberg adjusted, with the following thresholds:

- Alpha: 0.05
- Targeted power: >0.8 (met for primary compositional analyses)
- FDR cutoff: 0.05
- CV threshold: <100% (achieved here, ensuring biological plausibility).

### Differential Expression Analysis

Differential expression (DE) analysis was conducted to identify genes that are significantly up- or down-regulated between cell types or conditions. We utilized the estimateDEPerCellType function from the Cacoa R package^52^ (version 0.4.0; https://github.com/kharchenkolab/cacoa) which is specifically designed for differential analyses in sc/snRNA-seq experiments. Initially, counts were aggregated by sample within each cell type (pseudo-bulk, two biological replicates per region). These pseudo-bulk count matrices were then normalized using DESeq2’s median-of-ratios method, and low-count genes were removed by independent filtering (independent.filtering=TRUE). Differential expression between cell types was assessed by the DESeq2 Wald test (test = ‘DESeq2.Wald’, DESeq2 v1.42.0). Raw p-values were adjusted for multiple comparisons via the Benjamini–Hochberg procedure; genes with adjusted p-value (FDR < 0.05 were deemed significant. To ensure the accuracy and reliability of DE detection, we applied independent filtering using the Wald test. This approach ensures that only genes with sufficient expression and robust variance are considered for further analysis.

### Gene Ontology, Pathway Enrichment Analysis and Orthologs

To interpret the biological significance of differentially expressed genes, we performed over-representation analysis of Gene Ontology (GO) terms. For this, we used clusterProfiler (v4.10.0) with the org.Mmu.eg.db annotation package (v3.18.0) to test the Biological Process, Cellular Component, and Molecular Function categories; p-values were adjusted using the Benjamini– Hochberg method. Full results (including GO ID, term name, gene ratios, raw/adjusted p-values, gene IDs, gene counts, cell type and gene expression enrichment categories for each cell-type comparison) are provided in Supplementary Data 1 and visualized in Supplementary Fig. 5.

Protein sequence orthologs for NLGN1 (KEGG entry: hsa:22871), NTNG1 (hsa:22854), SPOCK1 (hsa:6695), RORB (hsa:6096), and KCNH8 (hsa:131096) were identified using the KEGG Sequence Similarity DataBase (SSDB; https://www.genome.jp/kegg/genes.html). SSDB contains precomputed pairwise similarities for protein-coding genes across KEGG genomes, generated via the SSEARCH program for amino acid sequences. For each human gene (Homo sapiens; code: hsa), best-hit orthologs (by similarity score) were retrieved for mouse (Mus musculus; mmu), common marmoset (Callithrix jacchus; cjc), and rhesus macaque (Macaca mulatta; mcc) using the SSDB “Best Hits” tool. Percentage identity (%ID) values, extracted directly from results, reflect identical amino acid residues over aligned sequence length ([identical residues / alignment length] × 100).

### In situ hybridization

In situ hybridization was performed according to the manufacturer’s instructions as stated in the protocol for fresh-frozen tissue (ACD – RNAScope Multiplex Fluorescent Reagent Kit v2 ((Cat No. 323100)). In brief, 10 µm sections were cut, fixed at 4 °C for 30 min, dehydrated and treated with Hydrogen Peroxidase and Protease IV. The following RNAScope probes for *Macaca mulatta* (Mmu) were purchased: Mmu-CAMK2A-C1 (Cat No. 461731), Mmu-RORB-C2 (Cat No. 876301-C2), Mmu-PDE1A-C3 (Cat No. 898751-C3), Mmu-PARD3-C3 (Cat No. 898731-C3), Mmu-CRISPLD1-C3 (Cat No. 898741-C3) and Mmu-ZNF385D-C3 (Cat No. 898711-C3). The probes were hybridized at 40 °C for 2 h, and the signal was amplified. The probes were visualized using the Opal-520 (visualizing C1), Opal-570 (visualizing C2) and Opal-690 (visualizing C3) (Akoya Cat No. NEL810001KT).

### Image analysis and quantification

Images spanning all layers of the cortex were acquired at the Microscopy Core Facility at Paul-Ehrlich-Institut, Langen, Germany, on a Leica SP8 confocal laser scanning microscope using the tiling mode. Image analysis was performed in QuPath^59^ (v0.3.0) using the Analysis Guidelines from ACD (MK 51-154) for Fluorescent RNAScope image analysis.

Cortical layers were determined with the help of DAPI and Nissl staining, CAMK2A and RORB visualization. Every layer was split in 5 roughly equal parts and marked with rectangles. Analysis was performed per rectangle. In brief, nuclei were detected, and outline was enlarged by 3 µm for assuming a cell. Within the cell, positive probes above a manually set threshold were detected and counted. Thresholds were set per channel (e.g., DAPI >50 intensity units; probe >3 spots/cell for positivity). A cell was defined as positive when more than 3 probes were detected. The percentage was calculated as cells positive for CAMK2A and RORB and target over cells being positive for CAMK2A and RORB. Statistical testing was performed using the chi-square test (alpha=0.05, power=0.8 for n=20 windows), and correction for multiple comparisons used the Bonferroni method.

## Data Availability

We have deposited raw snRNA-seq data in BioProject under accession PRJNA1304247. Processed datasets, including UMAP coordinates and marker lists, are available via interactive platforms

V1 dataset: http://kkh.bric.ku.dk/p2auto/index.html?fileURL=http:/rdo/project_1/visualisation_2.bin

V4 dataset: http://kkh.bric.ku.dk/p2auto/index.html?fileURL=http:/rdo/project_1/visualisation_3.bin

Integrated dataset: http://kkh.bric.ku.dk/p2auto/index.html?fileURL=http:/rdo/project_1/visualisation_1.bin).

Public datasets referenced for cross-validation (e.g., Allen Brain RNA-seq for macaque V1) are cited with accessions. All other data supporting findings are in supplements or from corresponding authors^23,24,60^.

## Acknowledgments

The authors would like to acknowledge Julien Vezoli (INSERM) for his advice on layer segmentation. The authors would like to thank Roland Plesker (Paul-Ehrlich-Institut) for extracting and providing two macaque brains and Irina Korshunova (BRIC) for her experimental support.

## Author contributions

Conceptualization: PF, KK

Methodology: DMG, MYB, RDO, VP, TW, PF

Investigation: DMG, MYB, RDO, VP

Visualization: DMG, MYB, RDO, VP

Supervision: CJB, PF, KK

Writing—original draft: DMG, MYB, RDO, PF, KK

Writing—review & editing: DMG, MYB, RDO, VP, TW, CJB, PF, KK

