## Supplementary Tables and Figures for "Single-Cell Atlas Uncovers Layer 4 Heterogeneity and Functional Gradients in Rhesus Macaque Visual Cortex"

##### Supplementary Table 1: Key Gene Orthologs Across Species.

| Gene | Human Ortholog (% ID) | Mouse Ortholog (KEGG: mmu - % ID) | Marmoset Ortholog (KEGG: cjc - % ID) | Macaque Ortholog (KEGG: mcc - % ID) | Function and complementary information's |
|---|---|---|---|---|---|
| ***NLGN1*** | 100% (self) | 95.7% | 97.4% | 99.3% | Synapse formation [32]; autism [33-35]<br><br>https://www.kegg.jp/ssdb-bin/ssdb_best?org_gene=hsa:22871 |
| ***NTNG1*** | 100% (self) | 96.5% | 99.4% | 100% | Neurite outgrowth [29]; schizophrenia [31]<br><br>https://www.kegg.jp/ssdb-bin/ssdb_best?org_gene=hsa:22854 |
| ***SPOCK1*** | 100% (self) | 95.2% | 96.8% | 99.1% | Neural development[36,37]<br><br>https://www.kegg.jp/ssdb-bin/ssdb_best?org_gene=hsa:6695 |
| ***RORB*** | 100% (self) | 97.9% | 99.6% | 100% | L3-5 high expression in visual profiling[24]<br><br>Layer marker; association metrics (r, p): ~2.0, p<1e-300 [23]<br><br>https://www.kegg.jp/ssdb-bin/ssdb_best?org_gene=hsa:6096 |
| ***KCNH8*** | 100% (self) | 92.2% | 97.8% | 99.2% | L4/5 IT enrichment in V1[23]<br><br>Ion channel; association metrics (r, p): ~2.0, p<1e-300 [23]<br><br>https://www.kegg.jp/ssdb-bin/ssdb_best?org_gene=hsa:131096 |

% ID from KEGG orthologous database (SSDB, see links in the last columns for more informations)

##### Supplementary Table 2: Layer-Specific Marker Percentages.

| Area | Layer 2/3 (%) | Layer 4 (%) | Layer 5 (%) | Notes |
|---|---|---|---|---|
| **V1** | 12.4 ± 2.1 | 25.3 ± 4.1 | 9.8 ± 1.7 | Highest in layer 4; p<0.05 vs. V4 |
| **V2** | 10.2 ± 1.9 | 18.7 ± 3.2 | 8.5 ± 1.5 | PDE1A peak (~35%) |
| **V4** | 7.6 ± 1.4 | 8.2 ± 2.3 | 6.9 ± 1.2 | Lower density |
| **MT** | 6.5 ± 1.3 | 7.1 ± 1.8 | 5.4 ± 1.0 | Dorsal decline |
| **TEO** | 9.3 ± 1.6 | 12.5 ± 2.5 | 7.2 ± 1.3 | Ventral increase |

Means ± 95% CI for PARD3+ subtype; similar for others in Supplementary Table S2; p<0.001 for V1 layer 4 enrichments.

**Supplementary Table 3: Rhesus Macaque Animal Metadata.**

| Animal | Sex | Birth Date | Treatment | Notes | Euthanasia Reason | Date of Death | Assay performed |
|---|---|---|---|---|---|---|---|
| **223** | m | 34846 | - | father of 354 | end of experiment | 2019-01-28 | snRNAseq |
| **354** | f | 39501 | 2x peudotyped LV injection, 1x into the lymph nodes, 1x IV | - | end of experiment | 2019-01-20 | snRNAseq |
| **E** | m | 38856 | implated with electrode | - | hypercalcemia of the heart | 2020-10-07 | smFISH |
| **S** | f | 37377 | implanted with chamber | - | mesenchymal neoplasia of the serous membranes in the abdominal cavity | 2020-06-26 | smFISH |

**Supplementary Figure 1. FANS gating, Sequencing Quality Metrics, and Sample Metadata for snRNA-seq in Rhesus Macaque Visual Areas V1 and V4.**

(A) FANS gating strategy. Nuclei were gated on FSC-A/SSC-A plot. Further, singlets (FSC-w/FSC-H and SSC-W/SSC-H plots) were selected with following gating for 7AAD+ nuclei and NeuN+ neuronal nuclei. (B) Number of genes per nucleus detected for each sample. Line inside the box represents median, lower and upper hinges of the box correspond to the first and third quartiles. Upper and lower whiskers correspond to the smallest and the largest values, and not more than 1.5 x inter-quartile range. Y-axis was square root transformed. (C) Number of single nuclei sequenced for each sample. (D) Fraction of cell per region for each sample (E) Table listing the 10x kit version, the region, the hemisphere, the sex and the animal ID per sample. (F) Fraction of nuclei per subtype shown for samples. Black line sets apart the fractions corresponding with tissue sampled from V1 (above black line) and V4 (below black line).

**Supplementary Figure 2. Extended snRNA-seq dataset.**

(A) Major neuronal families and their subtypes were detected in V1 and V4 using single nuclei RNA sequencing. UMAP representation of single nuclei transcriptomes is shown; nuclei are colored by their subtype.

**Supplementary Figure 3. Impact of covariates.**

UMAP plots showing the distribution for the main co-variates and batches of the samples on the common embedding for (A) animals F354 and M224, (B) sex, (C) brain regions V1 and V4, (D) the dissected hemispheres, (E) the 10x genomic kit versions and (F) sequencing runs on Illumina chips.

**Supplementary Figure 4. Extended compositional differences in cell types within visual areas.**

(A) Top plot: V1 neuronal subtypes (in color) segregate from V4 subtypes (in grey) mostly by the presence of V1-specific neuronal subtypes of L4_RORB family; Bottom plot: Similarly, V4 subtypes (in color) segregate from V1 subtypes (in grey) mostly by the presence of V4-specific subtype L4_RORB_KCNH8_ABCA12. UMAP representation of nuclei transcriptomes for all sub families are shown. (B) Expression of marker genes being selected for further FISH analysis based on their expression in cell clusters uniquely present in V1 (layer 4). UMAP representation of nuclei transcriptomes for all sub families are shown.

**Supplementary Figure 5. Enriched GO terms (downregulated) show no similarities between unique V1 and V4 subtypes.**

(A) Heat map of the top three enriched GO terms (down) for each annotated cell type, colored by $-\log_{10}$(adjusted p-value). GO term enrichment was calculated using a hypergeometric test and adjusted for multiple comparisons by the Benjamini–Hochberg procedure. Terms were hierarchically clustered (complete linkage on Euclidean distance). Cell types unique to V1 are in green, unique to V4 in orange, and layer 4 types shared between V1 and V4 in purple.

Supplemetary Figure 1

A

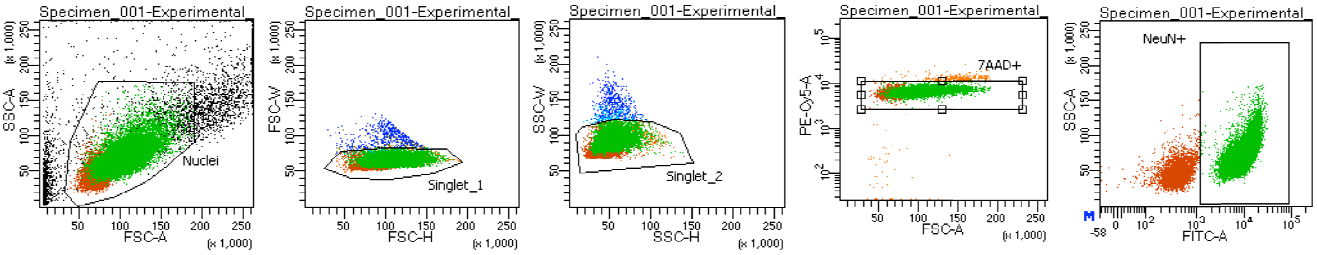


B

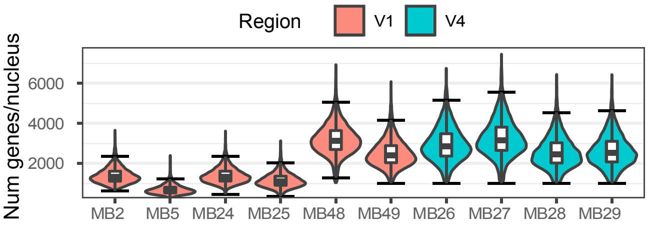


C

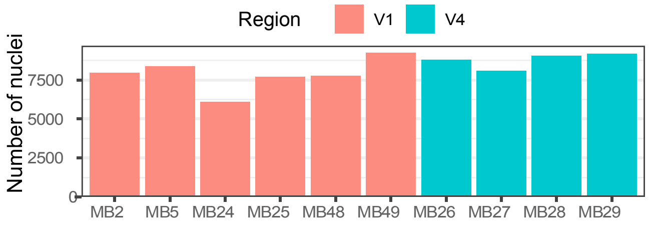


D

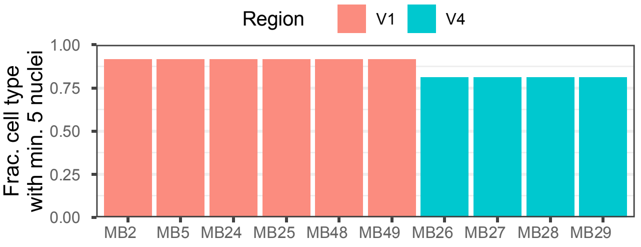


E

| Identifier | 10x kit version | Region | Hemisphere | Sex | Animal ID |
|---|---|---|---|---|---|
| MB2 | version 2 | V1 | right | F | F354 |
| MB5 | version 2 | V1 | left | M | M223 |
| MB24 | version 2 | V1 | left | M | M223 |
| MB25 | version 2 | V1 | right | F | F354 |
| MB48 | version 3 | V1 | left | M | M223 |
| MB49 | version 3 | V1 | left | F | F354 |
| MB26 | version 3 | V4 | left | F | F354 |
| MB27 | version 3 | V4 | left | F | F354 |
| MB28 | version 3 | V4 | left | M | M223 |
| MB29 | version 3 | V4 | left | M | M223 |

F

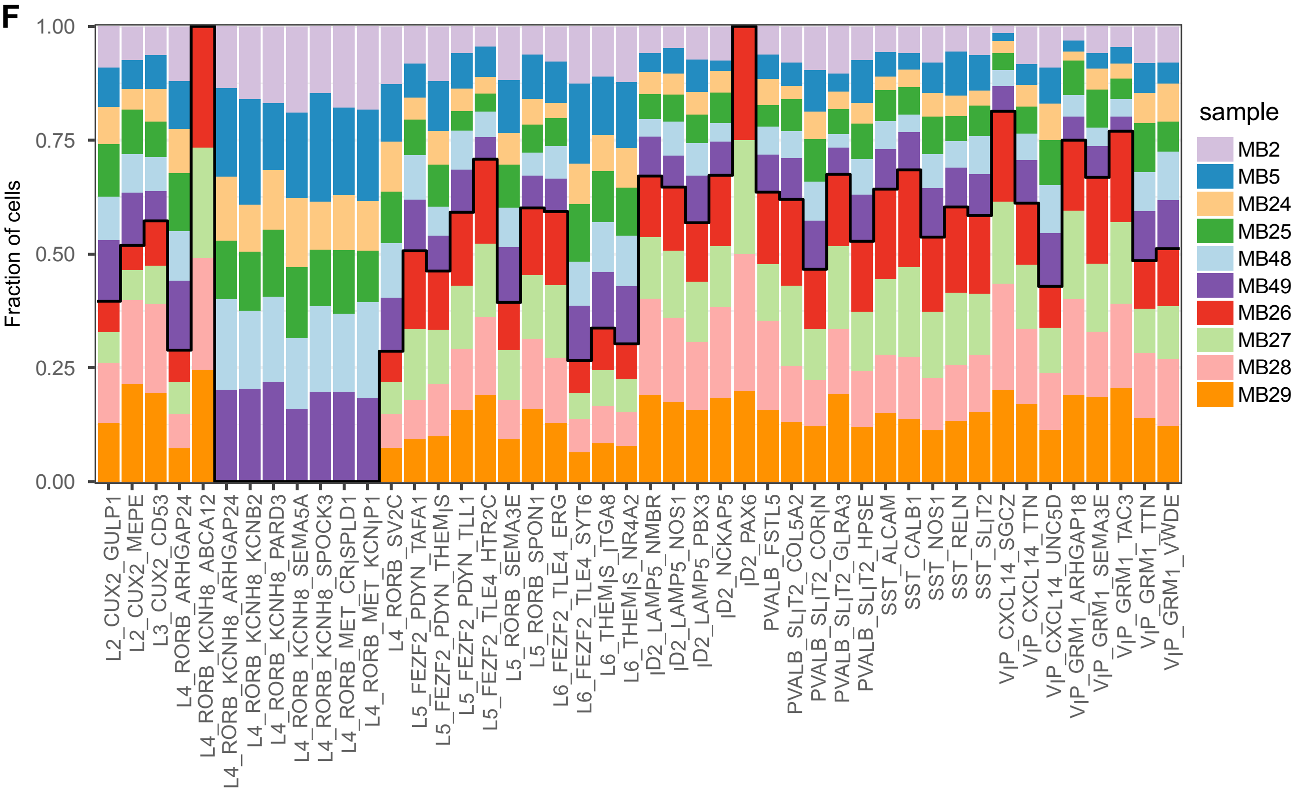

Supplementary Figure 2

**A**

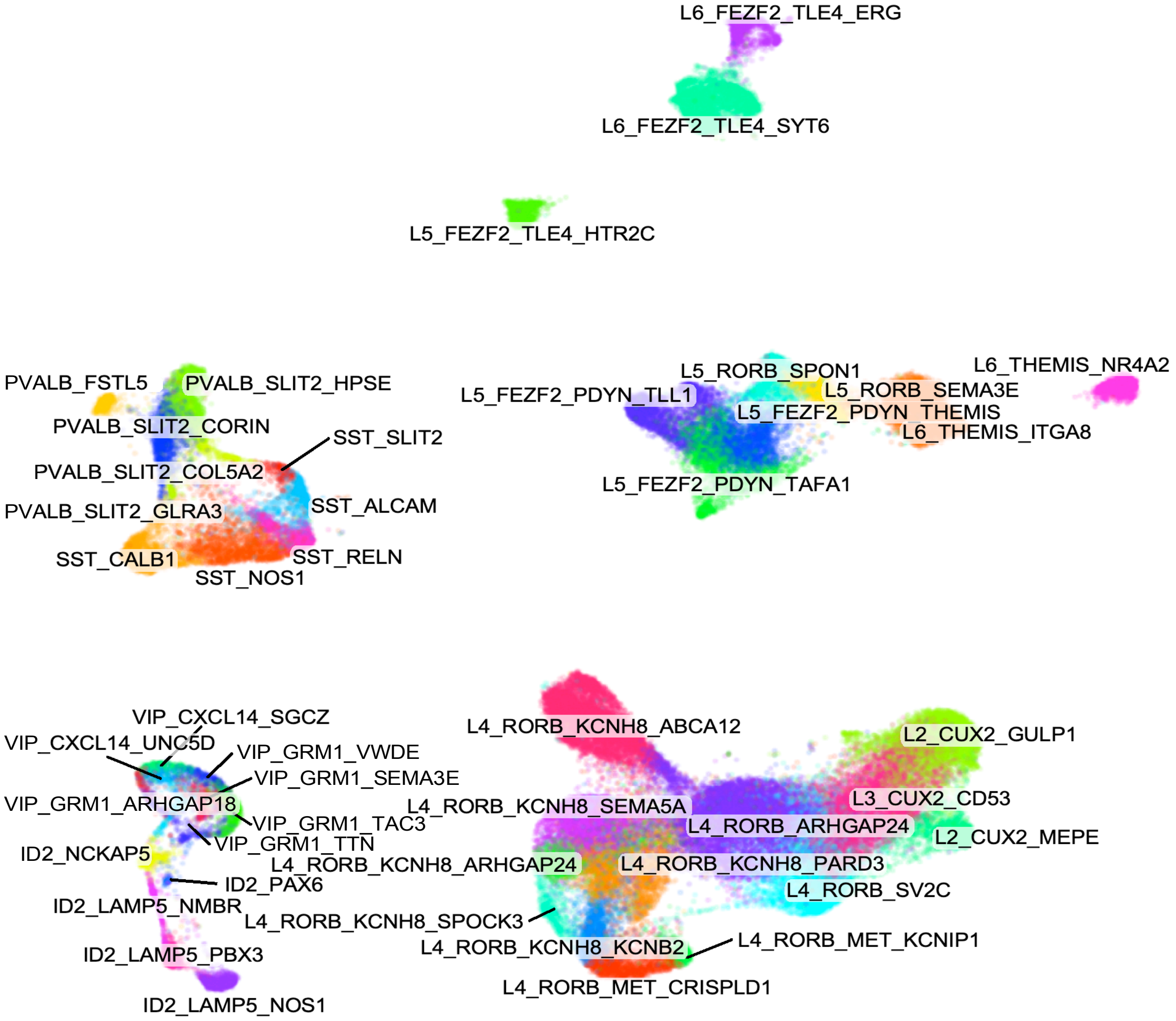

Supplementary Figure 3

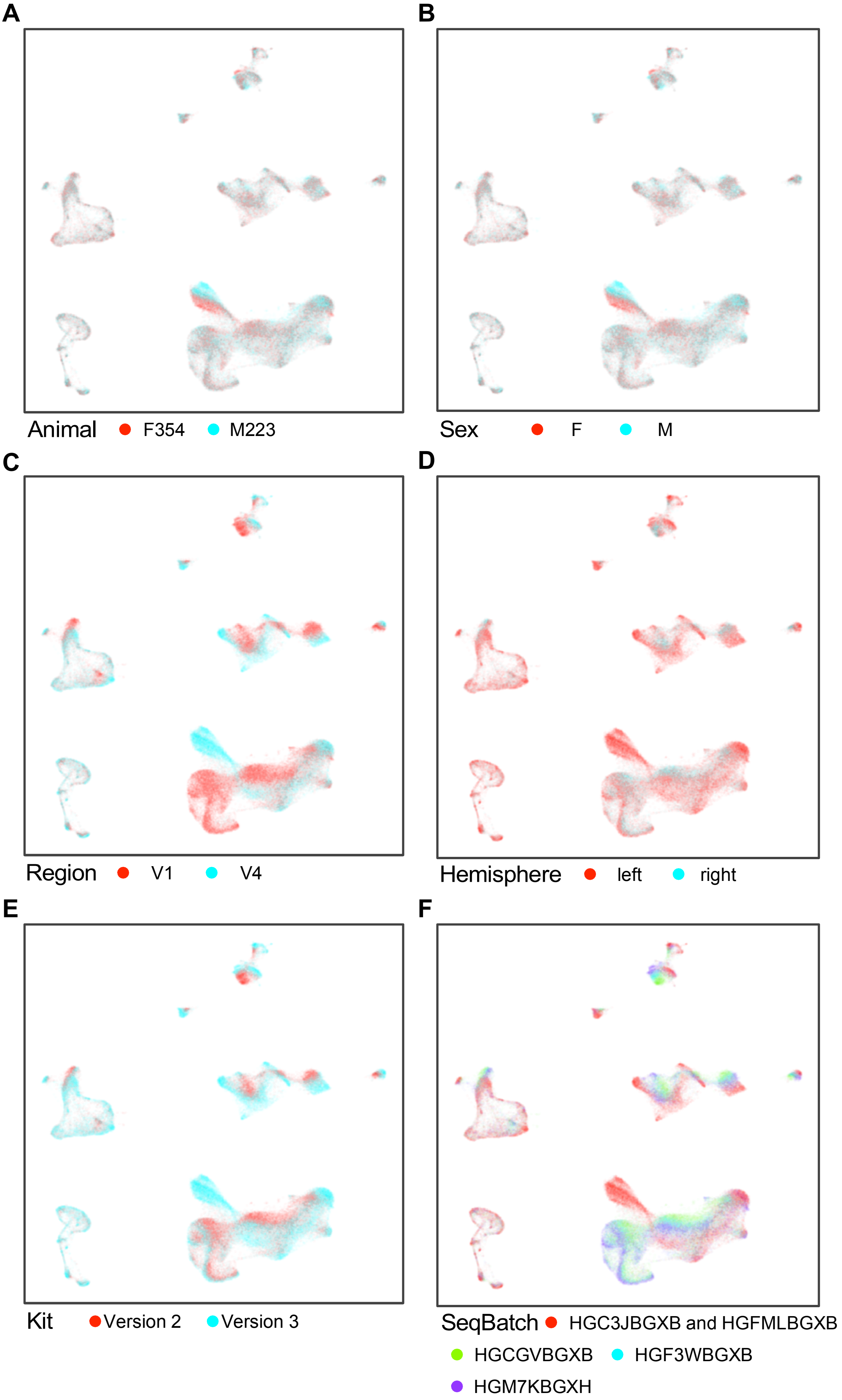

Supplementary Figure 4

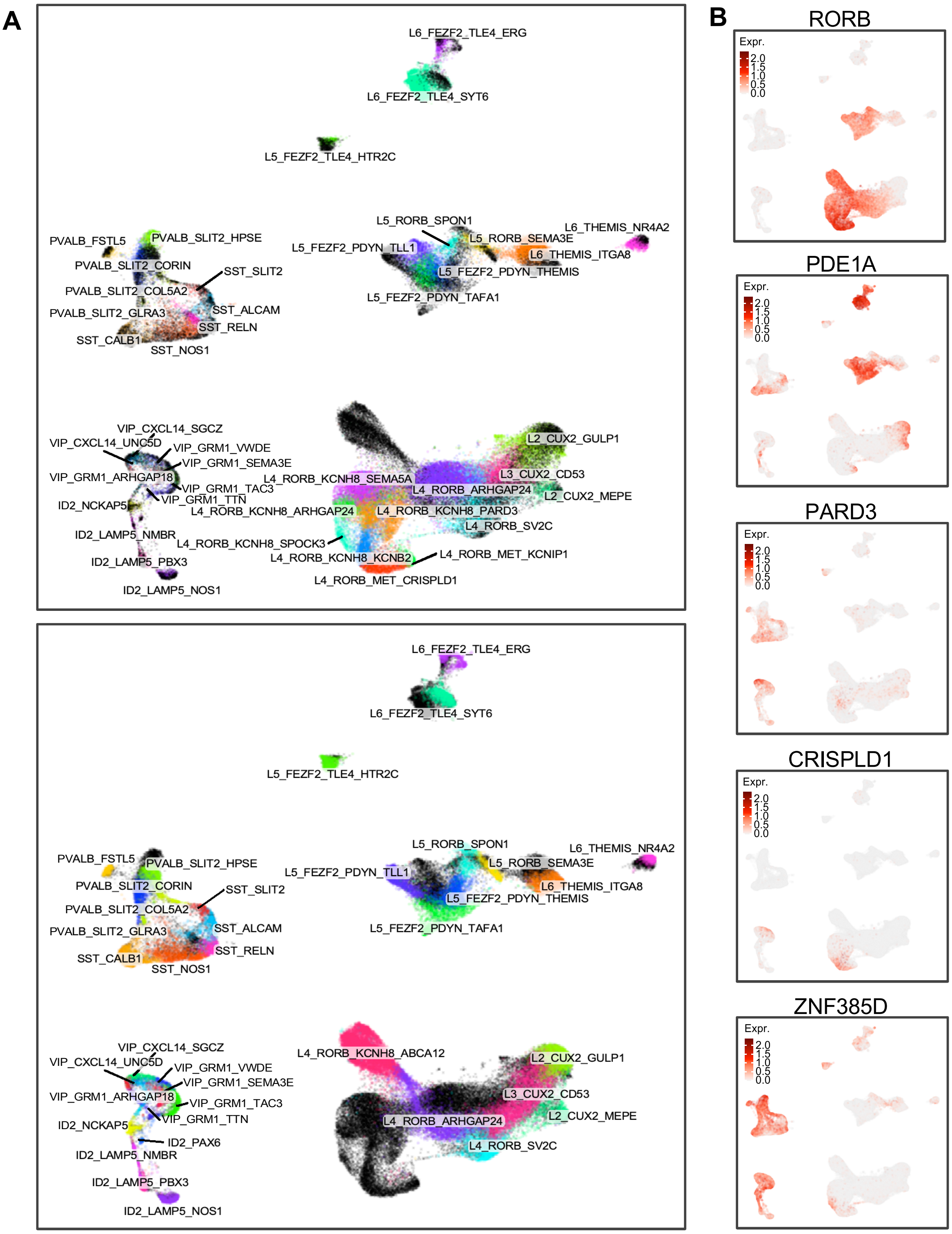

### Supplementary Figure 5

**A**

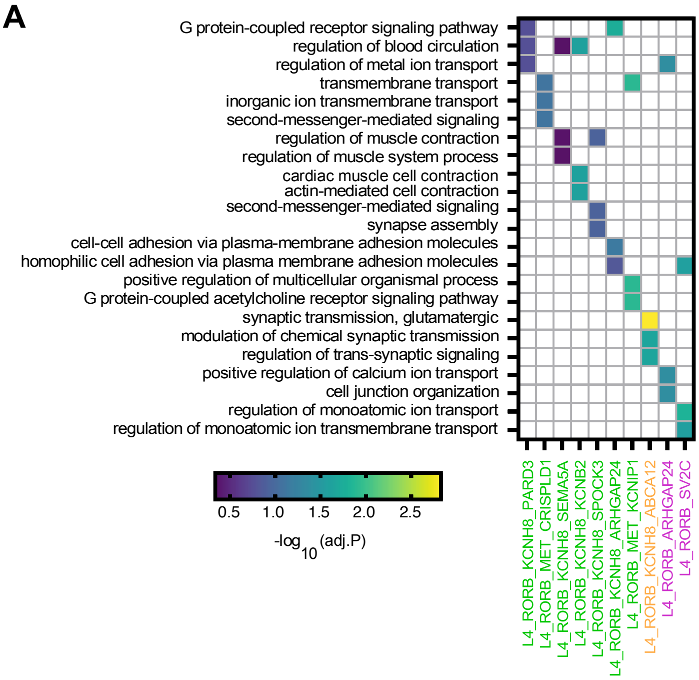
